# Omicron escapes the majority of existing SARS-CoV-2 neutralizing antibodies

**DOI:** 10.1101/2021.12.07.470392

**Authors:** Yunlong Cao, Jing Wang, Fanchong Jian, Tianhe Xiao, Weiliang Song, Ayijiang Yisimayi, Weijin Huang, Qianqian Li, Peng Wang, Ran An, Jing Wang, Yao Wang, Xiao Niu, Sijie Yang, Hui Liang, Haiyan Sun, Tao Li, Yuanling Yu, Qianqian Cui, Shuo Liu, Xiaodong Yang, Shuo Du, Zhiying Zhang, Xiaohua Hao, Fei Shao, Ronghua Jin, Xiangxi Wang, Junyu Xiao, Youchun Wang, Xiaoliang Sunney Xie

## Abstract

The SARS-CoV-2 B.1.1.529 variant (Omicron) contains 15 mutations on the receptor-binding domain (RBD). How Omicron would evade RBD neutralizing antibodies (NAbs) requires immediate investigation. Here, we used high-throughput yeast display screening^1,2^ to determine the RBD escaping mutation profiles for 247 human anti-RBD NAbs and showed that the NAbs could be unsupervised clustered into six epitope groups (A-F), which is highly concordant with knowledge-based structural classifications^3-5^. Strikingly, various single mutations of Omicron could impair NAbs of different epitope groups. Specifically, NAbs in Group A-D, whose epitope overlap with ACE2-binding motif, are largely escaped by K417N, G446S, E484A, and Q493R. Group E (S309 site)^6^ and F (CR3022 site)^7^ NAbs, which often exhibit broad sarbecovirus neutralizing activity, are less affected by Omicron, but still, a subset of NAbs are escaped by G339D, N440K, and S371L. Furthermore, Omicron pseudovirus neutralization showed that single mutation tolerating NAbs could also be escaped due to multiple synergetic mutations on their epitopes. In total, over 85% of the tested NAbs are escaped by Omicron. Regarding NAb drugs, the neutralization potency of LY-CoV016/LY-CoV555, REGN10933/REGN10987, AZD1061/AZD8895, and BRII-196 were greatly reduced by Omicron, while VIR-7831 and DXP-604 still function at reduced efficacy. Together, data suggest Omicron would cause significant humoral immune evasion, while NAbs targeting the sarbecovirus conserved region remain most effective. Our results offer instructions for developing NAb drugs and vaccines against Omicron and future variants.

## Main

The severe acute respiratory syndrome coronavirus 2 (SARS-CoV-2) variant B.1.1.529 was first reported to the World Health Organization (WHO) on 24 November 2021. It appears to be rapidly spreading, and the WHO classified it as a variant of concern (VOC) only two days after, designating it as Omicron ^8,9^. An unusually large number of mutations are found in Omicron, including over 30 in the spike protein (Extended Data Fig. 1a). The receptor-binding domain, responsible for interacting with the Angiotensin-Converting Enzyme 2 (ACE2) receptor, bears 15 of these mutations, including G339D, S371L, S373P, S375F, K417N, N440K, G446S, S477N, T478K, E484A, Q493R, G496S, Q498R, N501Y, and Y505H. Some of these mutations are very concerning due to their well-understood functional consequences, such as K417N and N501Y, which contribute to immune escape and higher infectivity ^10-13^. Many other mutations’ functional impacts remain to be investigated.

The S protein is the target of essentially all NAbs found in the convalescent sera or elicited by vaccines. Most of the N-terminal domain (NTD) neutralizing antibodies target an antigenic “supersite” in NTD, involving the N3 (residues 141 to 156) and N5 (residues 246 to 260) loops ^14,15^, and are thus very prone to NTD mutations. Omicron carries the Δ143-145 mutation, which would alter the N3 loop and most likely result in immune escape of most anti-NTD NAbs (Extended Data Fig. 1b). Compared to NTD targeting NAbs, RBD targeting NAbs are particularly abundant and potent, and display diverse epitopes. Evaluating how Omicron affects the neutralization capability of anti-RBD NAbs of diverse classes and epitopes is urgently needed.

RBD-directed SARS-CoV-2 NAbs can be assigned into different classes or binding sites based on structural analyses by cryo-EM or high-resolution crystallography ^3-5^; however, structural data only indicates the contacting amino acids, but does not infer the escaping mutations for a specific antibody. Recent advances in deep antigen mutation screening using FACS (fluorescence-activated cell sorting)-based yeast display platform has allowed the quick mapping of all single amino acid mutations in the RBD that affect the binding of SARS-CoV-2 RBD NAbs ^1,16^. The method has proven highly effective in predicting NAB drug efficacy toward mutations ^2^. However, to study how human humoral immunity may react to highly mutated variants like Omicron requires mutation profiling of a large collection of NAbs targeting different regions of RBD, and FACS-based yeast display mutation screening is limited by low experimental throughput. Here we further developed a MACS (magnetic-activated cell sorting) -based screening method which increases the throughput near 100-fold and could obtain comparable data quality like FACS (Fig 1a, Extended Data Fig. 2). Using this method, we quickly characterized the RBD escaping mutation profile for a total of 247 NAbs (Supplementary Data 1). Half of the NAbs were part of the antibodies identified by us using single-cell VDJ sequencing of antigen-specific memory B cells from SARS-CoV-2 convalescents, SARS-CoV-2 vaccinees, and SARS-CoV-1 convalescents who recently received SARS-CoV-2 vaccines (Supplementary Data 2). The other half of NAbs were identified by groups worldwide ^3,5,6,11,17-40^ (Supplementary Table 1).

**Fig. 1:**
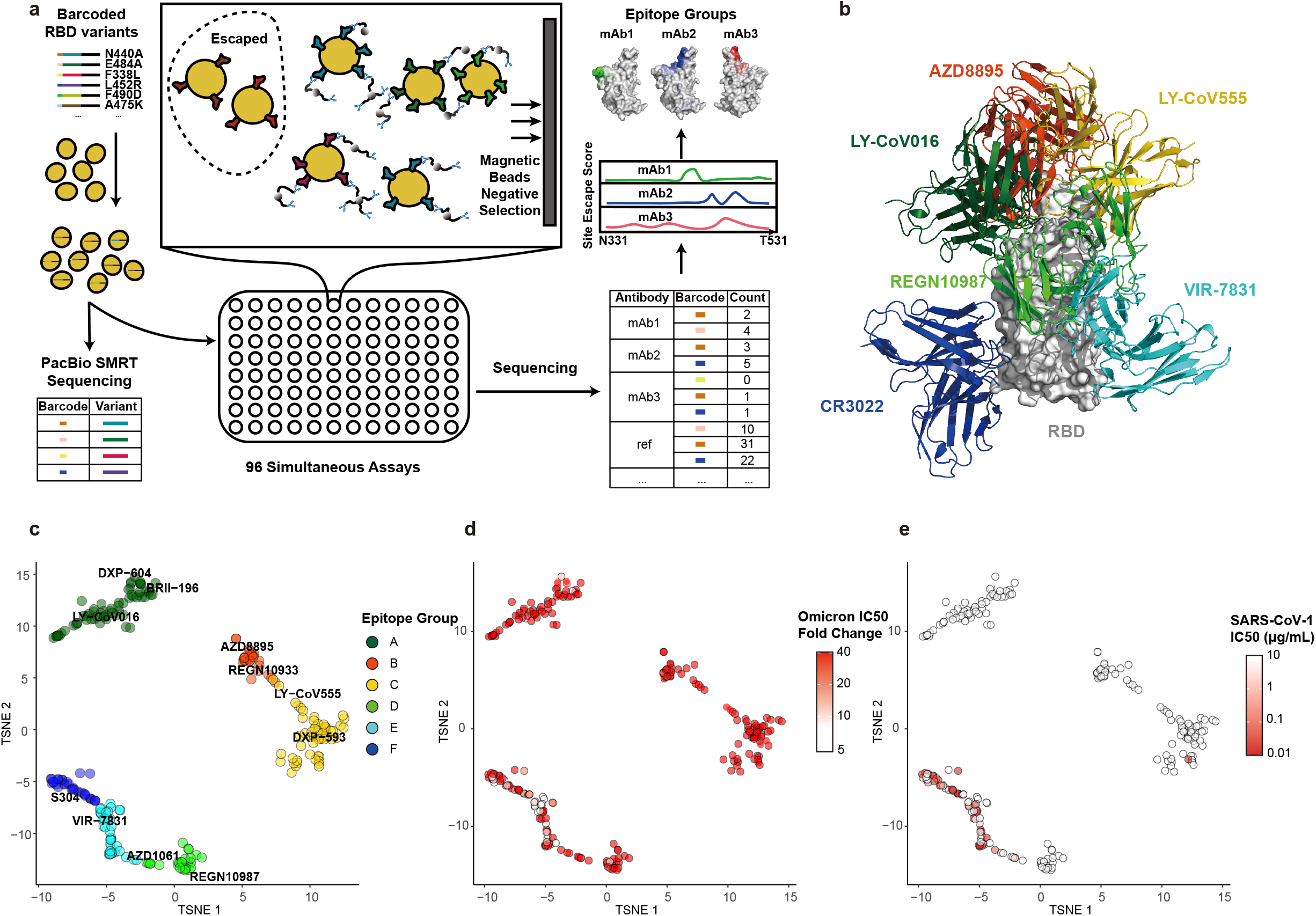
Omicron greatly reduces the neutralization potency of NAbs of diverse epitopes. **a**, Schematic of MACS-based high-throughput yeast display mutation screening. **b**, Representative NAb structures of each epitope group. **c**, t-SNE embedding and unsupervised clustering of SARS-CoV-2 human NAbs based on each antibody escaping mutation profile. A total of 6 epitope groups (Group A-F) could be defined. **d**, Neutralization of Omicron variant (spike-pseudotyped VSV) by 247 RBD NAbs. Shades of red show IC50 fold change compared with D614G of each NAb. **e**, Neutralization of SARS-CoV-1 (spike-pseudotyped VSV) by 247 RBD NAbs. Shades of red show the IC50 value (μg/mL) of each NAb. All pseudovirus neutralization assays are conducted in biological duplicates or triplicates.

The high-throughput screening capability allowed us to classify these NAbs into six Epitope Groups (A-F) using unsupervised clustering without dependence on structural studies, and the grouping is highly concordant with the knowledge-based structural classifications ^3-5^ (Fig. 1b, c). In particular, Group A-D NAbs largely correspond to the RBS A-D NAbs described by Yuan et al. ^4^ and overlap with the class 1-2 NAbs described by Barnes et al. ^3^ in general. The epitopes of these NAbs largely overlap with RBD residues involved in the binding to ACE2. Group A and B NAbs, represented by LY-CoV016 and AZD8895, respectively, usually can only bind to the ‘up’ RBD; whereas most of the Group C and D members, such as LY-CoV555 and REGN-10987, bind to RBDs regardless of their ‘up’ and ‘down’ conformations. Group E and F NAbs are very similar to the class 3 and 4 NAbs described by Barnes et al. ^3^ and target the S309/VIR-7831 site and CR3022 site, which could exhibit pan-sarbecovirus neutralization capacity (Fig 1e). Most of these NAbs neutralize SARS-CoV-2 using mechanisms other than directly interfering with ACE2 binding.

Inferred from the escaping mutation profiles, various single mutations of Omicron could impair NAbs of different epitope groups (Extended Data Fig. 3). Specifically, NAbs in Group A-D, whose epitope overlaps with ACE2-binding motif, are largely escaped by single mutations of K417N, G446S, E484A, and Q493R. Also, a subset of NAbs of Group E and F are escaped by single mutations of G339D, N440K, S371L, S375F. However, due to the extensive mutations accumulated on Omicron’s RBD, studying NAb’s response to Omicron only in the single mutation context is insufficient. Indeed, Omicron pseudovirus neutralization and spike enzyme-linked immunosorbent assay (ELISA) showed that single mutation tolerating NAbs could also be escaped by Omicron due to multiple synergetic mutations on their epitopes (Fig 1d, Extended Data Fig. 3). In total, over 85% of the tested human NAbs are escaped, suggesting that Omicron could cause significant humoral immune evasion and potential antigenic shifting.

It is crucial to analyze how each group of NAbs reacts to Omicron to instruct the development of NAb drugs and vaccines. Group A NAbs mainly contains the *VH3-53/VH3-66* germline gene-encoded antibodies, which are abundantly present in our current collection of SARS-CoV-2 neutralizing antibodies ^17,21,22,26,41-43^, including several antibodies that have obtained emergency use authorization (CB6/LY-CoV016) ^19^ or are currently being studied in clinical trials (P2C-1F11/BRII-196, BD-604/DXP-604) ^18,44^ (Fig. 2a, Extended Data Fig. 4a). Group A NAbs often exhibit less somatic mutations and shorter CDR3 length compared to other groups (Extended Data Fig. 5a, b). The epitopes of these antibodies extensively overlap with the binding site of ACE2 and are often evaded by RBD mutations on K417, D420, F456, A475, L455 sites (Fig 2d, Extended Data Fig. 6a,7a). Most NAbs in Group A were already escaped by B.1.351 (Beta) strain (Extended Data Fig. 5d), specifically by K417N (Extended Data Fig. 8a), due to a critical salt bridge interaction between Lys417 and a negatively charged residue in the antibody (Fig. 2g). The NAbs that survived Beta strain, such as BRII-196 and DXP-604, are insensitive to the K417N single site change but could also be heavily affected by the combination of K417N and other RBD mutations located on their epitopes, like S477N, Q493R, G496S, Q498R, N501Y, and Y505H of Omicron, causing lost or reduction of neutralization (Fig 2d; Extended Data Fig. 7a).

**Fig. 2:**
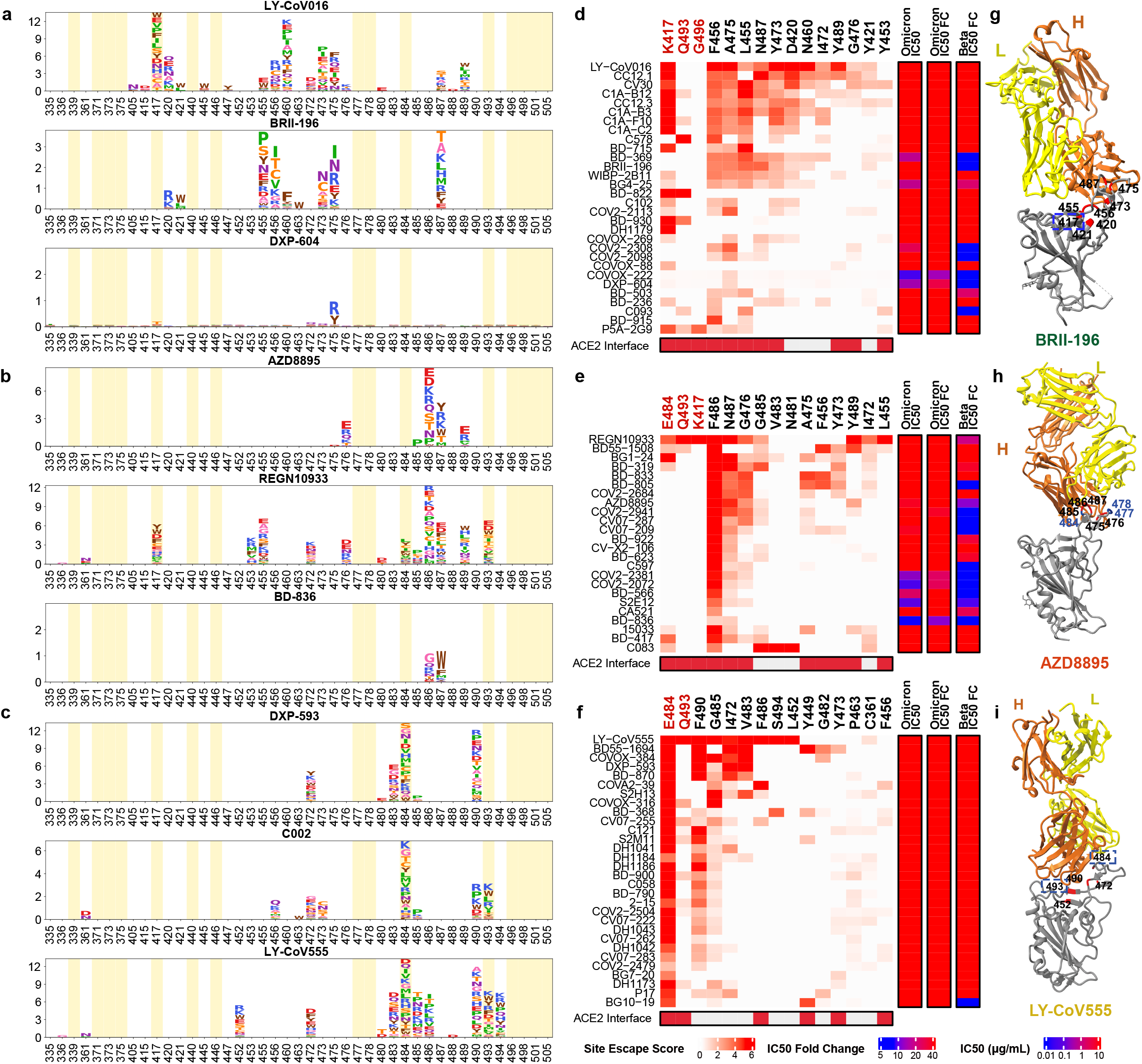
The neutralizing abilities of Group A-C NAbs are mostly abolished by Omicron. **a-c**, Escaping mutation profiles of representative NAbs for group A-C, respectively. For each site, the height of a letter indicates the detected mutation escape score of its corresponding residue. Sites mutated in Omicron are highlighted. **d-f**, Heatmaps of site escape scores for NAbs of epitope group A-C, respectively. ACE2 interface residues are annotated with red blocks, and mutated sites in Omicron are marked red. Annotations on the right side of heatmaps represent pseudovirus neutralizing IC50 fold change (FC) for Omicron and Beta compared to D614G. **g-i**, Representative structures of group A-C antibodies in complex with RBD. Residues involved in important contacts are labeled. Omicron mutations are marked as blue. NAb escaping mutations (Omicron) inferred from yeast display are labeled with squares.

The *VH1-58* gene-encoded NAbs are enriched in Group B (Extended Data Fig. 4b). These NAbs such as AZD8895 ^36^, REGN-10933 ^42^, and BD-836 ^45^ bind to the left shoulder of RBD, often focusing on the far tip (Fig. 2h). These NAbs are very sensitive to the change of F486, N487, and G476 (Fig 2b, Extended Data Fig. 6b). Fortunately, F486 and a few other major targeting sites of these NAbs are critically involved in ACE2-binding, and therefore they are generally harder to be escaped. A subset of NAbs in Group B, such as AZD8895 and BD-836, could survive Beta (Fig 2e); however, Omicron significantly reduced Group B NAbs’ binding affinity to RBD, potentially through S477N/T478K/E484A on their epitope (Extended Data Fig. 7b) ^46^, resulting in the loss of neutralization.

Group C NAbs are frequently encoded by *VH1-2* and *VH1-69* (Extended Data Fig. 4c). The majority of NAbs in this group could bind to both “up” and “down” RBDs, resulting in higher neutralization potency compared to other groups (Fig. 2c, Extended Data Fig. 5c). Several highly potent antibodies are found in Group C, including BD-368-2/DXP-593 ^44^, C002 ^3^, and LY-CoV555 ^47^. They bind to the right shoulder of RBD (Fig. 2i), and are mostly prone to the change of E484 (Extended Data Fig. 6c, 7c), such as the E484K mutation found in Beta (Fig. 2f). The E484A mutation seen in Omicron elicited a similar escaping effect, although the change to Ala is slightly subtler, and could be tolerated by certain antibodies in this group (Extended Data Fig. 8b). All Group C NAbs tested are escaped by Omicron.

Group D NAbs consist of diverse IGHV gene-encoded antibodies (Extended Data Fig. 4d). Prominent members in this group include REGN-10987 ^42^ and AZD1061 ^36^ (Fig. 3a). They further rotate down from the RBD right shoulder towards the S309 site when compared to Group C NAbs (Fig. 3g). As a loop formed by residues 440-449 in RBD is critical for the targeting of this group of NAbs, they are sensitive to the changes of N440, K444, G446, and N448 (Extended Data Fig. 6d, 7d). Most NAbs of Group D remain active against Beta; however, G446S would substantially affect their neutralization capability against Omicron (Fig. 3d). Also, for those NAbs that could tolerate G446S single mutation, the N440K/G446S combination may significantly reduce their binding affinity, resulting in that most Group D NAbs are escaped by Omicron.

**Fig. 3:**
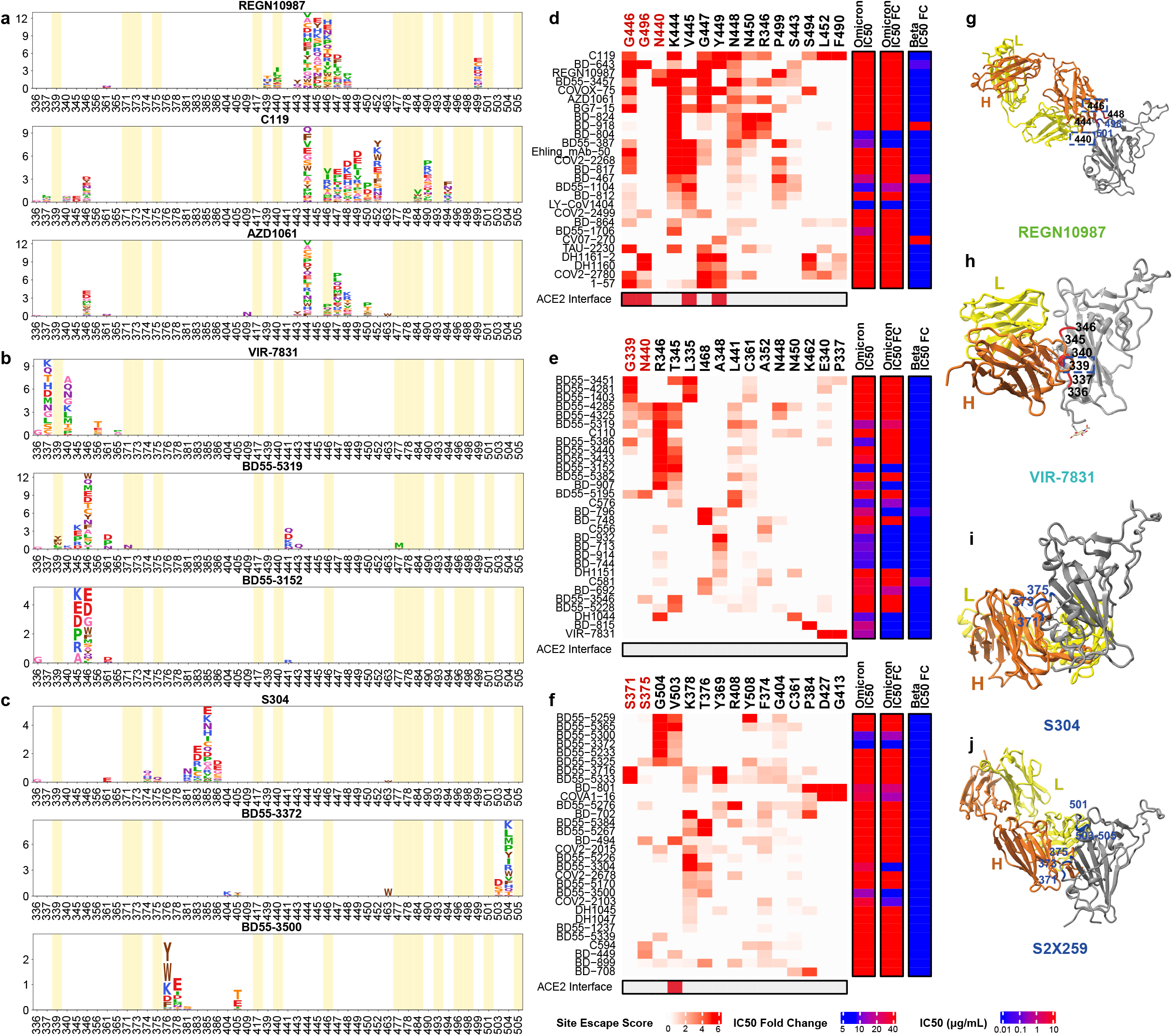
The majority of Group D-E NAbs are escaped by Omicron. **a-c**, Escaping mutation profiles of representative NAbs for group D-E, respectively. For each site, the height of a letter indicates the detected mutation escape score of its corresponding residue. Sites mutated in Omicron are highlighted. **d-f**, Heatmaps of site escape scores for NAbs of epitope group D-E, respectively. ACE2 interface residues are annotated with red blocks, and mutated sites in Omicron are marked red. Annotations on the right side of heatmaps represent pseudovirus neutralizing IC50 fold change (FC) for Omicron and Beta compared to D614G. **g-j**, Representative structures of group D-E antibodies in complex with RBD. Residues involved in important contacts are labeled. Omicron mutations are marked as blue. NAb escaping mutations (Omicron) inferred from yeast display are labeled with squares.

Group E and F NAbs are rarer when compared to the other four groups. The archetypical member of each group was originally isolated from a SARS-CoV-1 convalescent, and displays SARS-CoV-2 cross-neutralizing activity. There is no clear VDJ convergent effect compared to Group A, B, and C (Extended Data Fig. 4e, f), and the mutation rate and CDR3 length are larger than other groups. NAbs in Group E and F rarely compete with ACE2; thus, their average half-maximal inhibitory concentration (IC50) is higher than NAbs in Group A-D (Extended Data Fig. 5c). NAbs in Group E, such as VIR-7831/S309, may recognize a mixed protein/carbohydrate epitope, involving the N-linked glycan on N343 ^6^ (Fig. 3h). Inferred from the escaping mutation profiles (Fig. 3b), Group E NAbs are often sensitive to changes of G339, T345, and R346 (Extended Data Fig 6e, 7e). The G339D mutation would affect a subset of NAbs’ neutralization performance (Fig. 3e). Also, part of Group E NAbs’ epitope would extend to the 440-449 loop, making them sensitive to N440K in Omicron (Fig. 3e). Noticeably, the population of Omicron with R346K is continuously increasing, which may severely affect the neutralization capacity of Group E NAbs.

Group F NAbs such as S304 target a cryptic site in RBD that is generally not exposed (Fig. 3i), therefore their neutralizing activities are generally weaker ^7^. Group F NAbs are often sensitive to changes of F374, T376, and K378 (Extended Data Fig. 6f, 7f). A loop involving RBD residues 371-375 lies in the ridge between the E and F sites; therefore, a subset of Group F NAbs, including some Group E NAbs, could be affected by the S371L/S373P/S375F mutations if their epitopes extend to this region (Fig. 3c, f). Interestingly, a part of Group F NAbs is highly sensitive to V503 and G504, similar to the epitopes of S2X259 (Fig. 3f, j), suggesting that they can compete with ACE2. Indeed, several NAbs, such as BD55-5300 and BD55-3372, exhibit higher neutralization potency than other NAbs in Group F (Fig. 3c, 4b). However, These antibodies’ neutralization capability might be undermined by N501Y and Y505H of Omicron (Fig. 3j).

**Fig. 4:**
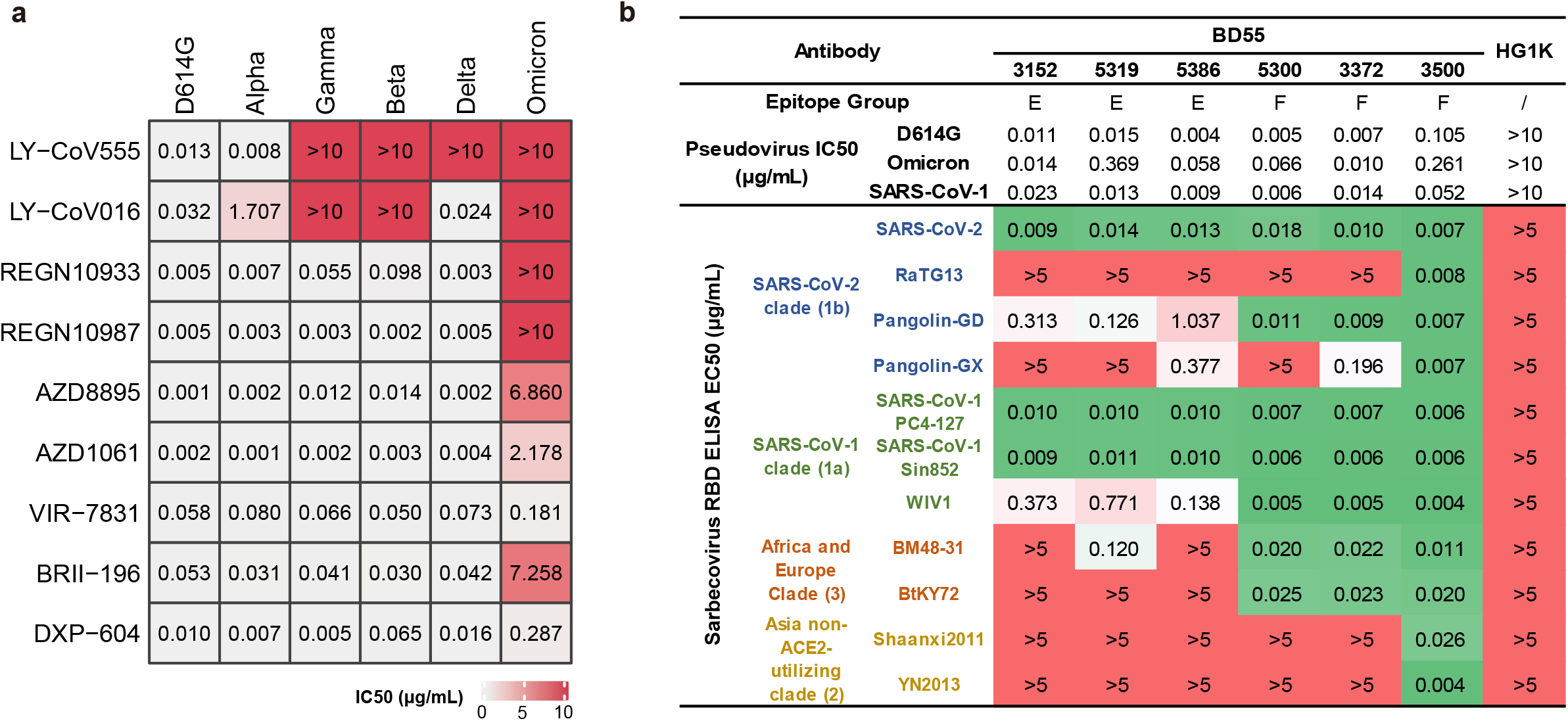
Omicron escapes most NAb drugs. **a**, Neutralization of SARS-CoV-2 variants of concern (pseudotyped VSV) by 9 NAb drugs. The pseudovirus neutralization assays for every VOC were performed in biological triplicates. IC50 labeled is the average of three replicates shown in Extended Data Fig. 9. **b**, The sarbecovirus neutralization and binding capability of selected potent Omicron-neutralizing antibodies. Monoclonal antibody HG1K (IgG1 antibody against Influenza A virus subtype H7N9) was used as the negative control.

As for NAb drugs, consistent with their escaping mutation profiles, the neutralization potency of LY-CoV016/LY-CoV555, REGN-10933/REGN-10987, and AZD1061 are greatly reduced by Omicron (Fig. 4a, Extended Data Fig. 9). The binding affinity of AZD8895 and BRII-196 toward Omicron RBD is also significantly reduced, likely due to multiple mutations accumulating on their epitopes, such that AZD8895 and BRII-196 failed to neutralize Omicron (Extended Data Fig. 10). BRII-198 was not tested since the antibody sequence was not released. VIR-7831 retains strong RBD binding capability, although G339 is part of its epitope, the G339D mutation in Omicron does not appear to affect VIR-7831’s binding; however, VIR-7831’s IC50 is reduced to 181 ng/mL, and may be subject to further reduction against Omicron with R346K. DXP-604’s binding affinity against Omicron RBD is largely reduced compared to wildtype RBD; nevertheless, it can still neutralize Omicron at an IC50 of 287 ng/mL, a nearly 30-fold reduction compared to wildtype (Fig. 4a). Additionally, several NAbs in Group E and F have shown high potency against Omicron and broad pan-sarbecovirus neutralization ability, promising for NAb drug development (Fig. 4b). Many more NAbs identified from vaccinated SARS-CoV-1 convalescents are waiting to be characterized.

The high-throughput yeast screening method provides a laboratory means for quickly examining the epitope of a certain NAb; however, the current throughput using FACS is limited and can not be used to evaluate a large NAb library. By virtue of MACS, we are able to increase the throughput by two orders of magnitude. In doing so, we were able to gain statistical confidence for the survival proportion of anti-RBD NAbs in each epitope group against Omicron. The experimental accuracy for predicting the neutralization reduction for single amino acid mutations is relatively high (Extended Data Fig. 8a, b); however, current mutation screening through yeast display could not effectively probe the consequence of multiple mutations simultaneously, which requires further technical optimization.

To date, a large number of SARS-CoV-2 anti-RBD NAbs have been identified from convalescents and vaccinees. The most potent NAbs are frequently found in Groups A-D as we described above, which tend to directly interfere with the binding of ACE2. Nevertheless, the neutralizing powers of these NAbs are often abrogated by RBD mutations in the evolutionary arms race between SARS-CoV-2 and human humoral immunity. Indeed, we showed that Omicron would escape the majority of SARS-CoV-2 NAbs in this collection (Extended Data Fig. 5e). On the other hand, Groups E and F NAbs are less affected by Omicron, likely because they are not abundant in population ^48^, hence exerting less evolutionary pressure for RBD to mutate in the corresponding epitope groups. These NAbs target conserved RBD regions in sarbecovirus and therefore are ideal targets for future development of pan-sarbecovirus NAb drugs.

## Methods

### Human peripheral blood mononuclear cells isolation

SARS-CoV-2 convalescents, SARS-CoV-1 convalescents, and SARS-CoV-2 vaccinees were recruited on the basis of prior SARS-CoV-2 infection or SARS-CoV-1 infection or SARS-CoV-2 at Beijing Youan and Ditan hospital. Relevant experiments regarding SARS-CoV-2 convalescents and vaccinees were approved by the Beijing Youan Hospital Research Ethics Committee (Ethics committee archiving No. LL-2020-010-K). Relevant experiments regarding SARS-CoV-1 convalescents were approved by the Beijing Ditan Hospital Capital Medical University (Ethics committee archiving No. LL-2021-024-02). All participants provided written informed consent for the collection of information, and that their clinical samples were stored and used for research. Data generated from the research were agreed to be published. The detailed information of SARS-CoV-2 convalescents and vaccinees was previously described ^11^. Briefly, short-term convalescents’ blood samples were obtained at day 62 on average after symptoms onset. Long-term convalescents’ blood samples were obtained at day 371 on average after symptoms onset. No vaccination was received before blood collection. SARS-CoV-2 vaccinees’ blood samples were obtained 2 weeks after complete vaccination of ZF2001 (RBD-subunit vaccine). For vaccinated SARS-CoV-1 convalescents (average age 58, n = 21), all recruited participants were previously identified for SARS-CoV-1 infection in 2003, and received two-dose vaccination of CoronaVac and a booster dose of ZF2001 with a 180-day-interval. 20mL of blood samples of the vaccinated SARS-CoV-1 convalescents were obtained 2 weeks after the booster shot. Three Healthy vaccinated donor (average age 25) were also included to serve as negative control for FACS gating. Peripheral Blood Mononuclear Cells (PBMCs) were separated from whole blood samples based on the detailed protocol described previously ^11^. Briefly, blood samples were first diluted with 2% Fetal Bovine Serum (FBS) (Gibco) in Phosphate Buffer Saline (PBS) (Invitrogen) and subjected to Ficoll (Cytiva) gradient centrifugation. After red blood cell lysis and washing steps, PBMCs were resuspended with 2% FBS in PBS for downstream B cell isolation or 10% Dimethyl sulfoxide (Sigma-Aldrich) in FBS for further preservation.

### Antigen-specific B cell sorting and sequencing

Starting with freshly isolated or thawed PBMCs, B cells were enriched by positive selection using a CD19+ B cell isolation kit according to the manufacturer’s instructions (STEMCELL). The enriched B cells were stained in FACS buffer (1× PBS, 2% FBS, 1 mM EDTA) with the following anti-human antibodies and antigens: For every 10^6 cells, 3 μL FITC anti-CD19 Antibody (Biolegend, 392508), 3 μL FITC anti-CD20 Antibody (Biolegend, 302304), 3.5 μL Brilliant Violet 421 anti-CD27 Antibody (Biolegend, 302824), 3 μL PE/Cyanine7 anti-IgM(Biolegend, 314532), and fluorophore-labelled Receptor-Binding Domain (RBD) and ovalbumin (Ova) for 30 min on ice. Cells were stained with 5 μL 7-AAD (eBioscience, 00-6993-50) for 10 minutes before sorting. Biotinylated receptor binding domain (RBD) of SARS (Sino biological, 40634-V27H-B) or SARS-CoV-2 (Sino biological, 40592-V27H-B) were multimerized with fluorescently labeled Streptavidin (SA) for 1 hour at 4°C. RBD was mixed with SA-PE (Biolegend, 405204) and SA-APC (Biolegend, 405207) at a 4:1 molar ratio. For every 10^6^ cells, 6 ng SA was used to stain. Single CD19 or CD20+, CD27+, IgM-, Ova-, RBD-PE+, RBD-APC+, live B cells were sorted on an Astrios EQ (BeckMan Coulter) into PBS containing 30% FBS (Supplementary Data 2). FACS sorting were controlled by Summit 6.0 (Beckman Coulter). FACS data analyses were done by FlowJo 10.8. Cells obtained after FACS were sent for 5′-mRNA and V(D)J libraries preparation as previously described^11^, which were further submitted to Illumina sequencing on a Hiseq 2500 platform, with the 26×91 pair-end reading mode.

### V(D)J sequence data analysis

The raw FASTQ files were processed by Cell Ranger (version 6.1.1) pipeline using GRCh38 reference. Sequences were generated using “cellranger multi” or “cellranger vdj” with default parameters. Antibody sequences were processed by IMGT/DomainGapAlign (version 4.10.2) to obtain the annotations of V(D)J, regions of complementarity determining regions (CDR), and the mutation frequency^49,50^. Mutation count divided by the length of the V gene peptide is defined as the amino acid mutation rate of the V gene.

### Recombinant antibody production

Paired immunoglobulin heavy and light chain genes obtained from 10X Genomics V(D)J sequencing and analysis were submitted to recombinant monoclonal antibody synthesis. Briefly, heavy and light genes were cloned into expression vectors, respectively, based on Gibson assembly, and subsequently co-transfected into HEK293F cells (ThermoFisher, R79007). The secreted monoclonal antibodies from cultured cells were purified by protein A affinity chromatography. The specificities of these antibodies were determined by ELISA.

### ELISA

ELISA plates were coated with RBD (SARS-CoV-2 WT, SARS-CoV-2 Omicron, SARS-CoV-1 RBD, Sino Biological Inc.) at 0.03 μg/mL and 1 μg/mL in PBS at 4°C overnight. After standard washing and blocking, 100 μL 1μg/mL antibodies were added to each well. After a 2 h incubation at room temperature, plates were washed and incubated with 0.08 μg/mL goat anti-human IgG (H+L)/HRP (JACKSON, 109-035-003) for 1 h incubation at room temperature. Tetramethylbenzidine (TMB) (Solarbio) was then added, and the reaction was stopped by adding H_2_SO_4_. OD450 was measured by an ELISA microplate reader. An antibody is defined as ELISA-positive when the OD450 (1 μg/mL RBD) is three times larger than the negative control, which utilizes an H7N9 specific human IgG1 antibody (HG1K, Sino Biology Cat #HG1K).

### Peudovirus neutralization assay

Pesudovisurs neutralization assay was performed to evaluate neutralizing ability of antibodies. The detailed process was previously described by Cao et al.^12^. Briefly, serially diluted antibodies were first incubated with pseudotyped virus for 1h, and the mixture was then incubated with Huh-7 cells. After 24h incubation in an incubator at 37°C, cells were collected and lysed with luciferase substrate (PerkinElmer), then proceeded to luminescence intensity measurement by a microplate reader. IC50 was determined by a four-parameter non-linear regression model using PRISM (v9.0.1). Omicron pseudovirus contains the following mutations: A67V, H69del, V70del, T95I, G142D, V143del, Y144del, Y145del, N211del, L212I, ins214EPE, G339D, S371L, S373P, S375F, K417N, N440K, G446S, S477N, T478K, E484A, Q493R, G496S, Q498R, N501Y, Y505H, T547K, D614G, H655Y, N679K, P681H, N764K, D796Y, N856K, Q954H, N969K, L981F.

### Biolayer interferometry

Biolayer interferometry assays were conducted on Octet® R8 Protein Analysis System (Fortebio) following the manufacturer’s instruction. Briefly, after baseline calibration, Protein A biosensors (Fortebio) were immersed with antibodies to capture the antibody, then sensors were immersed in PBS with 0.05% Tween-20 to the baseline. After association with different concentrations of RBD of SARS-CoV-2 variants (Omicron RBD: 40592-V08H85), disassociation was conducted. Data were recorded using Octet BLI Discovery (12.2) and analyzed using Octet BLI Analysis (12.2).

### RBD Deep Mutational Scanning Library construction

The yeast-display RBD mutant libraries used here were constructed as described by Starr et al.,^12^ based on the spike receptor-binding domain (RBD) from SARS-CoV-2 (NCBI GenBank: MN908947, residues N331-T531) with the modifications that instead of 16-neuclotide barcode (N16), a unique 26-neuclotide (N26), barcode was appended to each RBD variant as an identifier in order to decrease sequencing cost by eliminating the use of PhiX. Briefly, three rounds of mutagenesis PCR were performed with designed and synthesized mutagenetic primer pools; in order to solid our conclusion, we constructed two RBD mutant libraries independently. RBD mutant libraries were then cloned into pETcon 2649 vector and the assembled products were electroporated into electrocompetent DH10B cells to enlarge plasmid yield. Plasmid extracted form E. coli were transformed into the EBY100 strain of *Saccharomyces cerevisiae* via the method described by Gietz and Schiestl^51^. Transformed yeast population were screened on SD-CAA selective plate and further cultured in SD-CAA liquid medium at a large scale. The resulted yeast libraries were flash frozen by liquid nitrogen and preserved at -80°C.

### PacBio library preparation, sequencing, and analysis

The correspondence of RBD gene sequence in mutant library and N26 barcode was obtained by PacBio sequencing. Firstly, the bacterially-extracted plasmid pools were digested by NotI restriction enzyme and purified by agarose gel electrophoresis, then proceed to SMRTbell ligation. Four RBD mutant libraries were sequenced in one SMRT cell on a PacBio Sequel ll platform. PacBio SMRT sequencing subreads were converted to HiFi ccs reads with pbccs, and then processed with a slightly modified version of the script previously described^12^ to generate the barcode-variant dictionary. To reduce noise, variants containing stop codons or supported by only one ccs read were removed from the dictionary and ignored during further analysis.

### Magnetic-activated cell sorting (MACS)-based mutation escape profiling

ACE2 binding mutants were sorted based on magnetic beads to eliminate non-functional RBD variants. Briefly, the biotin binder beads (Thermo Fisher) were washed and prepared as the manufacturer’s instruction and incubated with biotinylated ACE2 protein (Sino Biological Inc.) at room temperature with mild rotation. The ACE2 bound beads were washed twice and resuspend with 0.1% BSA buffer (PBS supplemented with 0.1% bovine serum albumin), and ready for ACE2 positive selection. Transformed yeast library were inoculated into SD-CAA and grown at 30°C with shaking for 16-18h, then back-diluted into SG-CAA at 23°C with shaking to induce RBD surface expression. Yeasts were collected and washed twice with 0.1% BSA buffer and incubated with aforementioned ACE2 bound beads at room temperature for 30min with mild rotating. Then, the bead-bound cells were washed, resuspend with SD-CAA media, and grown at 30°C with shaking. After overnight growth, the bead-unbound yeasts were separated with a magnet and cultured in a large scale. The above ACE2 positive selected yeast libraries were preserved at -80°C in aliquots as a seed bank for antibody escape mapping.

One aliquot of ACE2 positive selected RBD library was thawed and inoculated into SD-CAA, then grown at 30°C with shaking for 16-18h. 120 OD units were back-diluted into SG-CAA media and induced for RBD surface expression. Two rounds of sequential negative selection to sort yeast cells that escape Protein A conjugated antibody binding were performed according to the manufacturer’s protocol. Briefly, Protein A magnetic beads (Thermo Fisher) were washed and resuspend in PBST (PBS with 0.02% Tween-20). Then beads were incubated with neutralizing antibody and rotated at room temperature for 30min. The antibody-conjugated beads were washed and resuspend in PBST. Induced yeast libraries were washed and incubated with antibody-conjugated beads for 30min at room temperature with agitation. The supernatant was separated and proceed to a second round of negative selection to ensure full depletion of antibody-binding yeast.

To eliminate yeast that did not express RBD, MYC-tag based RBD positive selection was conducted according to the manufacturer’s protocol. First, anti-c-Myc magnetic beads (Thermo Fisher) were washed and resuspend with 1X TBST (TBS with Tween-20), then the prepared beads were incubated for 30min with the antibody escaping yeasts after two rounds of negative selection. Yeasts bound by anti-c-Myc magnetic beads were wash with 1X TBST and grown overnight in SD-CAA to expand yeast population prior to plasmid extraction.

Overnight cultures of MACS sorted antibody-escaped and ACE2 preselected yeast populations were proceed to yeast plasmid extraction kit (Zymo Research). PCRs were performed to amplify the N26 barcode sequences as previously described^13^. The PCR products were purified with 0.9X Ampure XP beads (Beckman Coulter) and submitted to 75bp single-end Illumina Nextseq 500 sequencing.

### Deep mutational scanning data processing

Raw single-end Illumina sequencing reads were trimmed and aligned to the reference barcode-variant dictionary generated as described above to get the count of each variant with *dms_variants* Python package (version 0.8.9). For libraries with N26 barcodes, we slightly modified the *illuminabarcodeparser* class of this package to tolerate one low sequencing quality base in the barcode region. The escape score of variant X is defined as F×(n_X,ab_ / N_ab_) / (n_X,ref_ / N_ref_), where n_X,ab_ and n_X,ref_ is the number of detected barcodes for variant X, N_ab_ and N_ref_ are the total number of barcodes in antibody-selected (ab) library and reference (ref) library respectively as described by Starr et al. ^12^. Different from FACS experiments, as we couldn’t measure the number of cells retained after MACS selection precisely, here F is considered as a scaling factor to transform raw escape fraction ratios to 0-1 range, and is calculated from the first and 99th percentiles of raw escape fraction ratios. Scores less than the first percentile or larger than the 99th percentile are considered to be outliers and set to zero or one, respectively. For each experiment, barcodes detected by <6 reads in the reference library were removed to reduce the impact of sampling noise, and variants with ACE2 binding below -2.35 or RBD expression below -1 were removed as previously described ^12^. Finally, we built global epistasis models with dms_variants package for each library to estimate single mutation escape scores, utilizing the Python scripts provided by Greaney et al. ^16^. To reduce experiment noise, sites are retained for further analysis only if its total escape score is at least 0.01, and at least 3 times greater than the median score of all sites. For antibodies measured by 2 independent experiments, only sites which pass the filter in both experiments are retained. Logo plots in Fig. 2, Fig. 3, Extended Data Fig. 2 and Supplementary Data 1 are generated by Python package *logomaker* (version 0.8).

### Antibody clustering

Antibody clustering and epitope group identification were performed based on the N×M escape score matrix, where N is the number of antibodies which pass the quality controlling filters, and M is the number of informative sites on SARS-CoV-2 RBD. Each entry of the matrix A_nm_ refers to the total escape score of all kinds of mutations on site m of antibody n. The dissimilarity between two antibodies is defined based on the Pearson’s correlation coefficient of their escape score vectors, i. e. D_ij_=1-Corr(**A**_i_,**A**_j_), where Corr(**A**_i_, **A**_j_)=**x**_i_·**x**_j_/|**x**_i_||**x**_j_| and vector **x**_i_=**A**_i_-Mean(**A**_i_). Sites with at least 6 escaped antibodies (site escape score >1) were considered informative and selected for dimensionality reduction and clustering. We utilized R function *cmdscale* to convert the cleaned escape matrix into an N×6 feature matrix by multidimensional scaling (MDS) with the dissimilarity metric described above, followed by unsupervised k-medoids clustering within this 6-dimensional antibody feature space, using *pam* function of R package *cluster* (version 2.1.1). Finally, two-dimensional t-Distributed Stochastic Neighbor Embedding (tSNE) embeddings were generated with *Rtsne* package (version 0.15) for visualization. 2D t-SNE plots are generated by *ggplot2* (version 3.3.3), and heatmaps are generated by *ComplexHeatmap* package (version 2.6.2).

## Acknowledgments

We thank Professor Jesse Bloom for his generous gift of the yeast SARS-CoV-2 RBD libraries. We thank Beijing BerryGenomics for the help on DNA sequencing. We thank Sino Biological Inc. for the technical assistance on mAbs and Omicron RBD expression. We thank Sartorius (Shanghai) Trading Co., Ltd. for providing instrumental help with BLI measurement. We thank Jia Luo and Hongxia Lv (National Center for Protein Sciences and core facilities at School of Life Sciences at Peking University) for the help in flow cytometry. This project is financially supported by the Ministry of Science and Technology of China (CPL-1233).

## Author contributions

Y.C. and X.S.X designed the study. Y.C. and F.S coordinated the characterizations of the NAbs. J.W., F.J., H.L., H.S. performed and analyzed the yeast display mutation screening experiments. T.X., W.J., X.Y., P.W., H.L. performed the pseudovirus neutralization assays. W.H., Q.L., T.L., Y.Y., Q.C., S.L., Y.W. prepared the VSV-based SARS-CoV-2 pseudovirus. A.Y., Y.W., S.Y., R.A., W.S. performed and analyzed the antigen-specific single B cell VDJ sequencing. X.N., R.A. performed the antibody BLI studies. Z.C., S.D., P.L., L.W., Z.Z., X.W., J.X. performed the antibody structural analyses. P.W., Y.W., J.W, H.S, H.L. performed the ELISA experiments. X.H. and R.J. coordinated the blood samples of vaccinated SARS-CoV-1 convalescents. Y.C., X.W., J.X., X.S.X wrote the manuscript with inputs from all authors.

## Corresponding authors

Correspondence to Yunlong Cao or Xiangxi Wang or Junyu Xiao or Youchun Wang or Xiaoliang Sunney Xie. Request for materials described in this study should be directed to Xiaoliang Sunney Xie.

## Data availability

Data availabilityProcessed escape maps for NAbs are available in Supplementary Data 1 (as figures), or at https://github.com/sunneyxielab/SARS-CoV-2-RBD-Abs-HTDMS (as mutation escape score data). Raw Illumina and PacBio sequencing data are available on NCBI Sequence Read Archive BioProject PRJNA787091. We used vdj_GRCh38_alts_ensembl-5.0.0 as the reference of V(D)J alignment, which can be obtained from https://support.10xgenomics.com/single-cell-vdj/software/downloads/latest. IMGT/DomainGapAlign is based on the built-in lastest IMGT antibody database, and we let the “Species” parameter as “Homo sapiens” while kept the others as default. FACS-based deep mutational scanning datasets could be downloaded from https://media.githubusercontent.com/media/jbloomlab/SARS2_RBD_Ab_escape_maps/main/processed_data/escape_data.csv.

Processed data of this study has been added to this repository as well.

## Code availability

Scripts for analyzing SARS-CoV-2 escaping mutation profile data and for reproducing figures in this paper are available at https://github.com/sunneyxielab/SARS-CoV-2-RBD-Abs-HTDMS.

**Extended Data Fig. 1:**
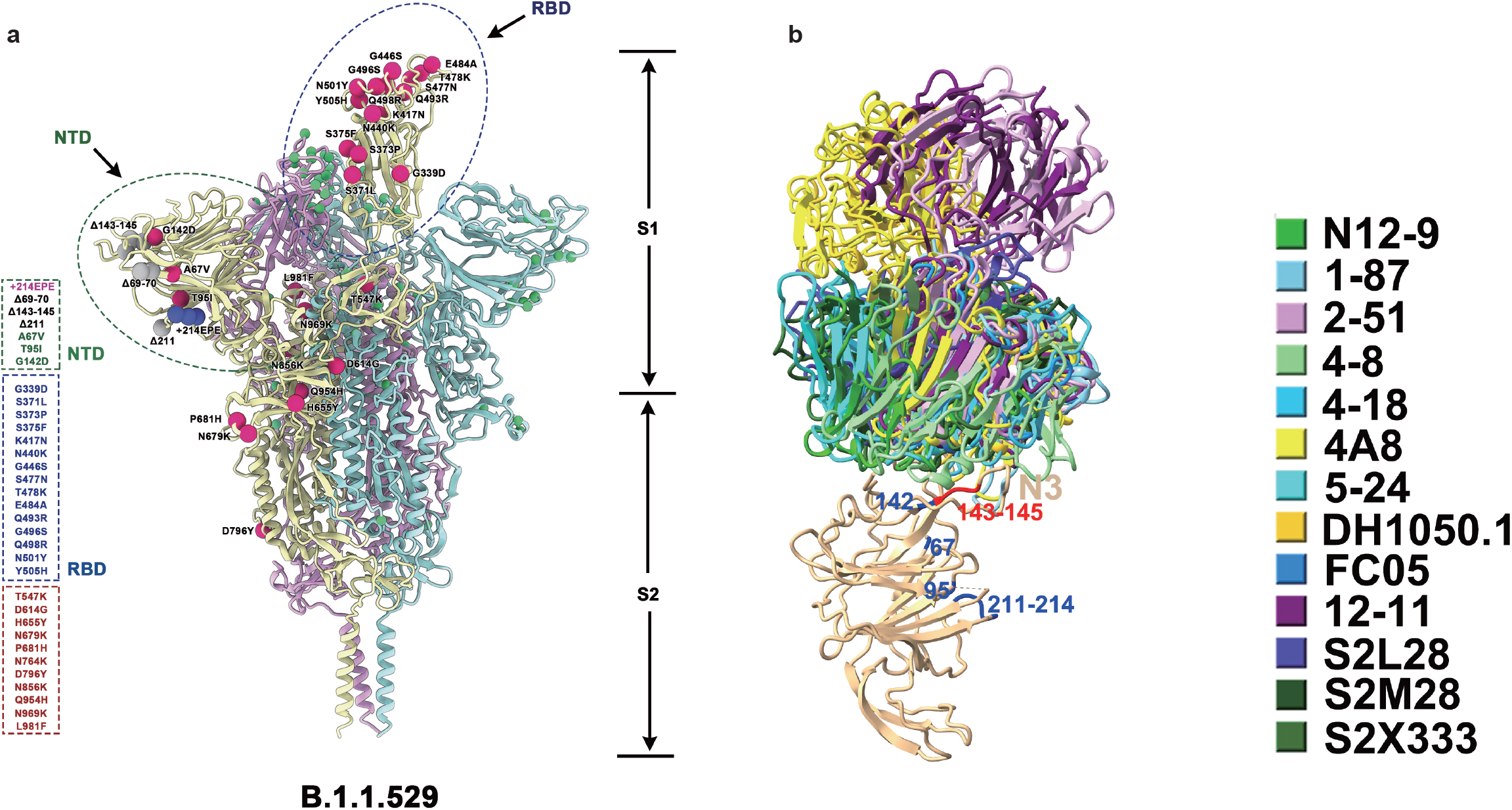
Illustration of SARS-CoV-2 spike with Omicron’s mutations. **a**, SARS-CoV-2 D614G spike protein structure overlayed with Omicron mutations. Omicron’s (BA.1) popular mutations are marked by red (for substitutions), blue (for insertions) and gray balls (for deletions). **b**, NTD-binding NAbs shown together in complex with NTD. Substitutions and deletions of Omicron NTD are colored blue and red, respectively.

**Extended Data Fig. 2:**
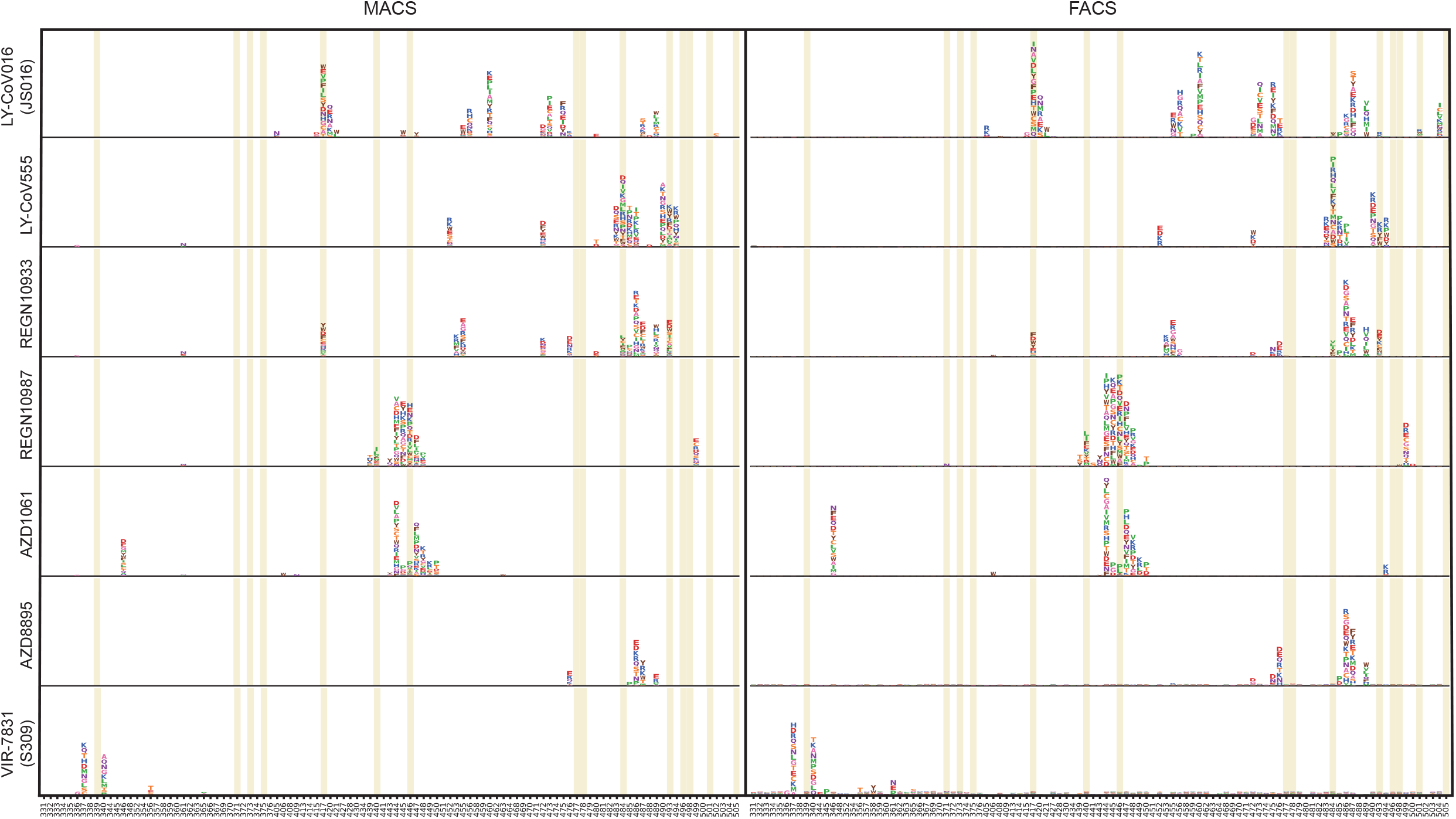
Comparison between FACS and MACS-based deep mutational scanning. Deep mutational scanning maps with MACS-based (left) and FACS-based assays (right) of seven therapeutic neutralizing antibodies that have received emergency use authorization. Sites mutated in the Omicron variant are highlighted. Mutation amino acids of each site are shown by single letters. The heights represent mutation escape score, and colors represent chemical properties. FACS-based data were obtained from public datasets by Jesse Bloom.

**Extended Data Fig. 3:**
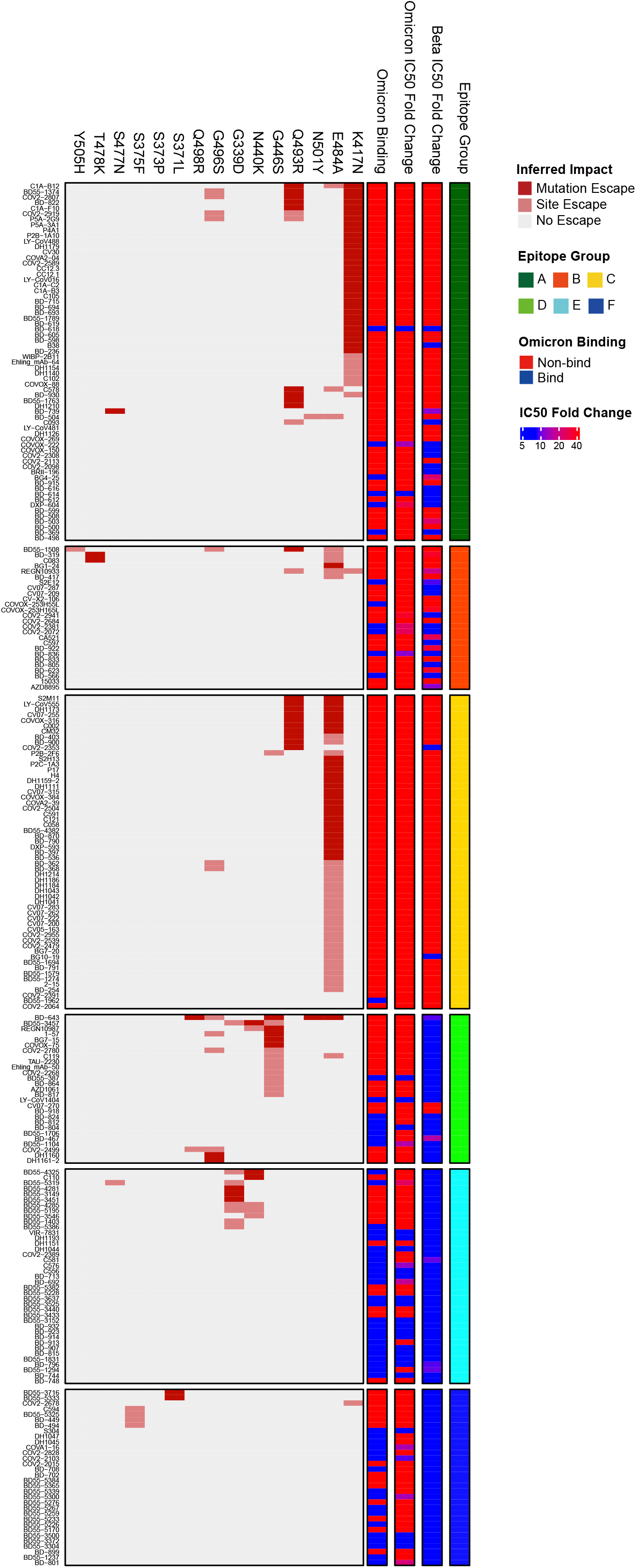
Omicron neutralization IC50 fold-change distribution of 247 NAbs of diverse epitopes. Fold-change of IC50 (VSV pseudovirus neutralization) compared to D614G by Beta and Omicron (BA.1) are shown for all 247 NAbs tested. The impact of each RBD mutation of Omicron on NAbs’ binding is inferred from yeast display mutation screening. Each NAb’s binding to Omicron RBD was validated through ELISA. All neutralization and ELISA assays were conducted in biological duplicates.

**Extended Data Fig. 4:**
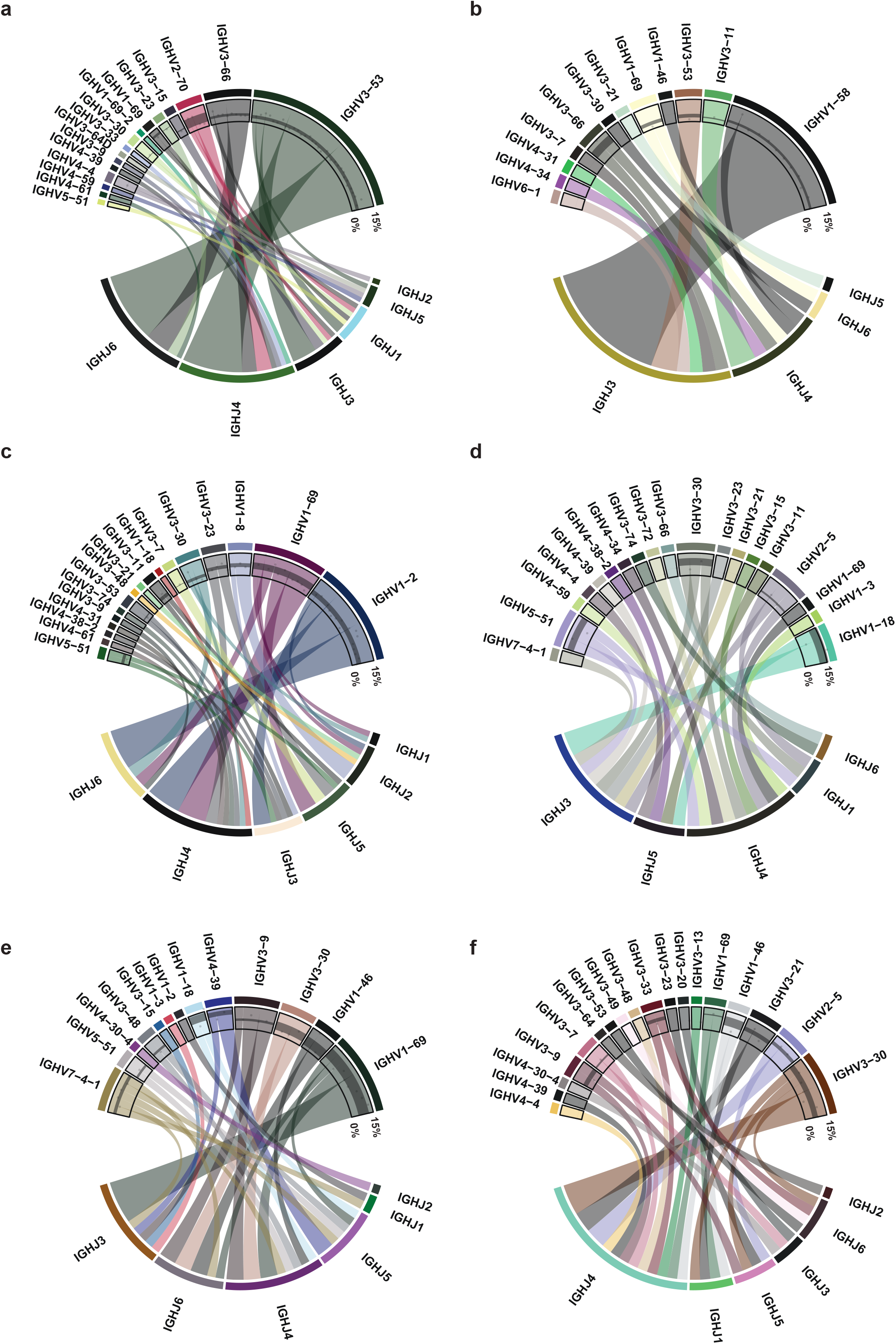
Heavy chain V/J segment recombination of NAbs of each epitope group. **a-f**, Chord diagrams showing the heavy chain V segment and J segment recombination of epitope group A(a), B(b), C(c), D(d), E(e) and F(f). The width of the arc linking a V segment to a J segment indicates the antibody number of the corresponding recombination. The inner layer scatter plots show the V segment amino acid mutation rate, while black strips show the 25%∼75% quantile of mutation rates.

**Extended Data Fig. 5.**
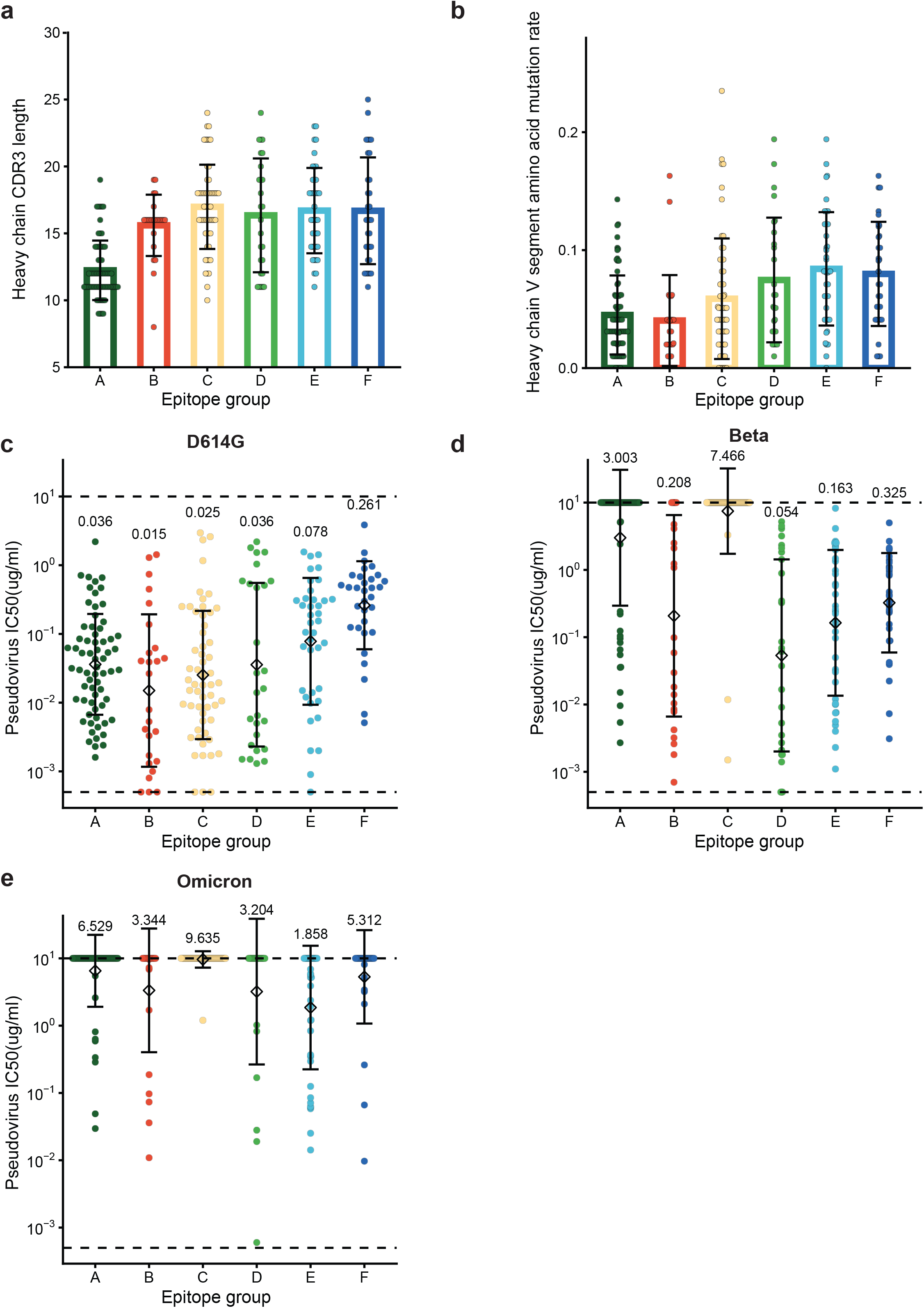
Neutralization potency, heavy chain CDR3 length, and mutation rate distribution for NAbs of each epitope group. **a**, The length of H chain complementarity-determining region 3 (HCDR3) amino acid sequence for NAbs in each epitope group (n=66, 26, 57, 27, 39, 32 antibodies for epitope group A, B, C, D, E, F, respectively). HCDR3 lengths are displayed as mean ± s.d. **b**, The V segment amino acid mutation rate for NAbs in each epitope group (n=66, 26, 57, 27, 39, 32 antibodies for epitope group A, B, C, D, E, F, respectively). Mutation rates are calculated are displayed as mean ± s.d. **c-e**, The IC50 against D614G(c), Beta(d), and Omicron(e) variants for NAbs in each epitope group (n=66, 26, 57, 27, 39, 32 antibodies for epitope group A, B, C, D, E, F, respectively). IC50 values are displayed as mean ± s.d. in the log10 scale. Pseudovirus assays for each variant are biologically replicated twice. Dotted lines show the detection limit, which is from 0.0005 μg/mL to 10 μg/mL. IC50 geometric means are also labeled on the figure.

**Extended Data Fig. 6:**
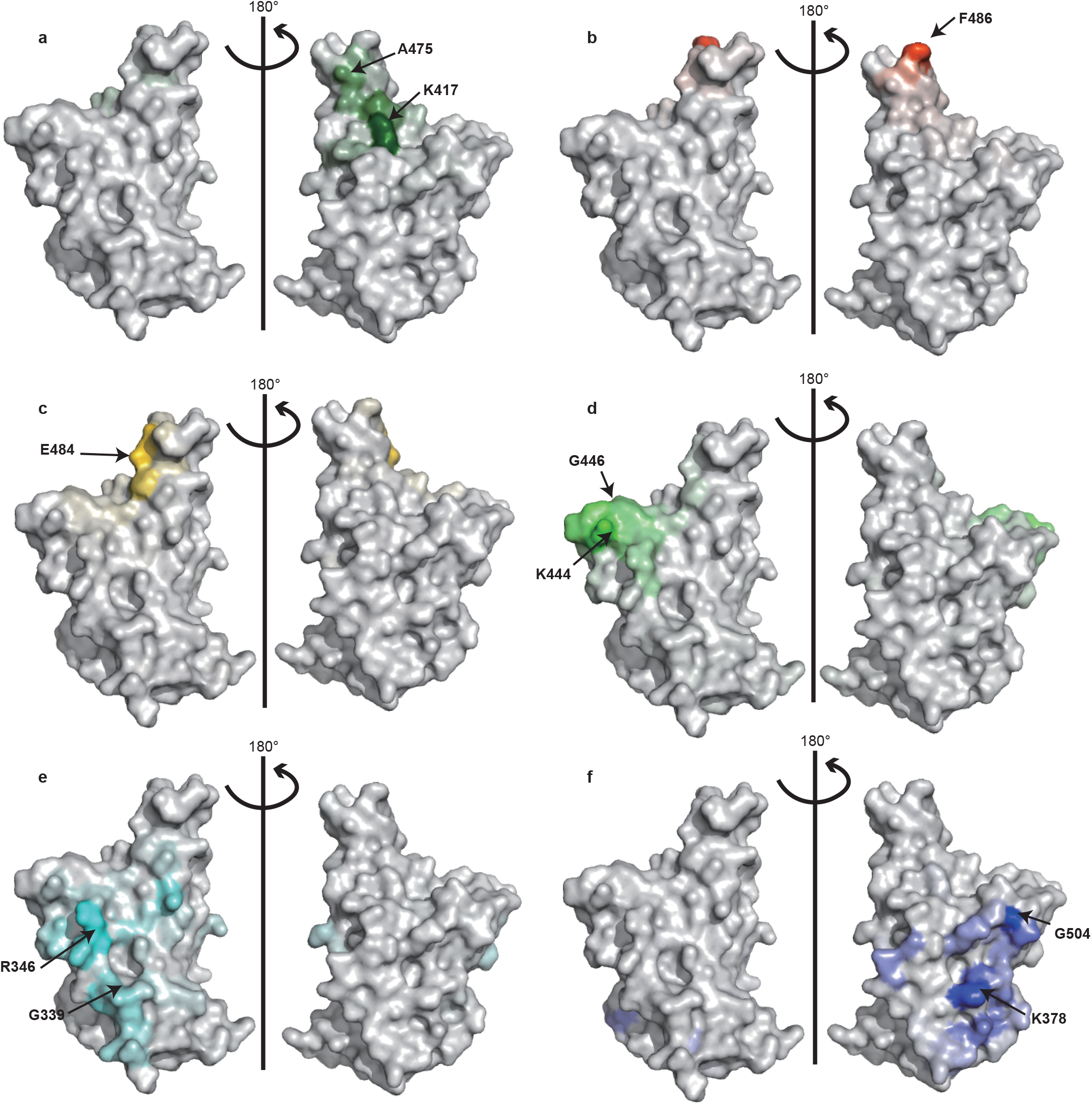
Escape hotspots of different epitope groups on the RBD surface. **a-f**, Aggregated site escape scores of antibodies for epitope group A-F, respectively. Epitope groups are distinguished by distinct colors, and the shades show normalized site escape scores. Escape hotspots of each epitope group are annotated by arrows.

**Extended Data Fig. 7:**
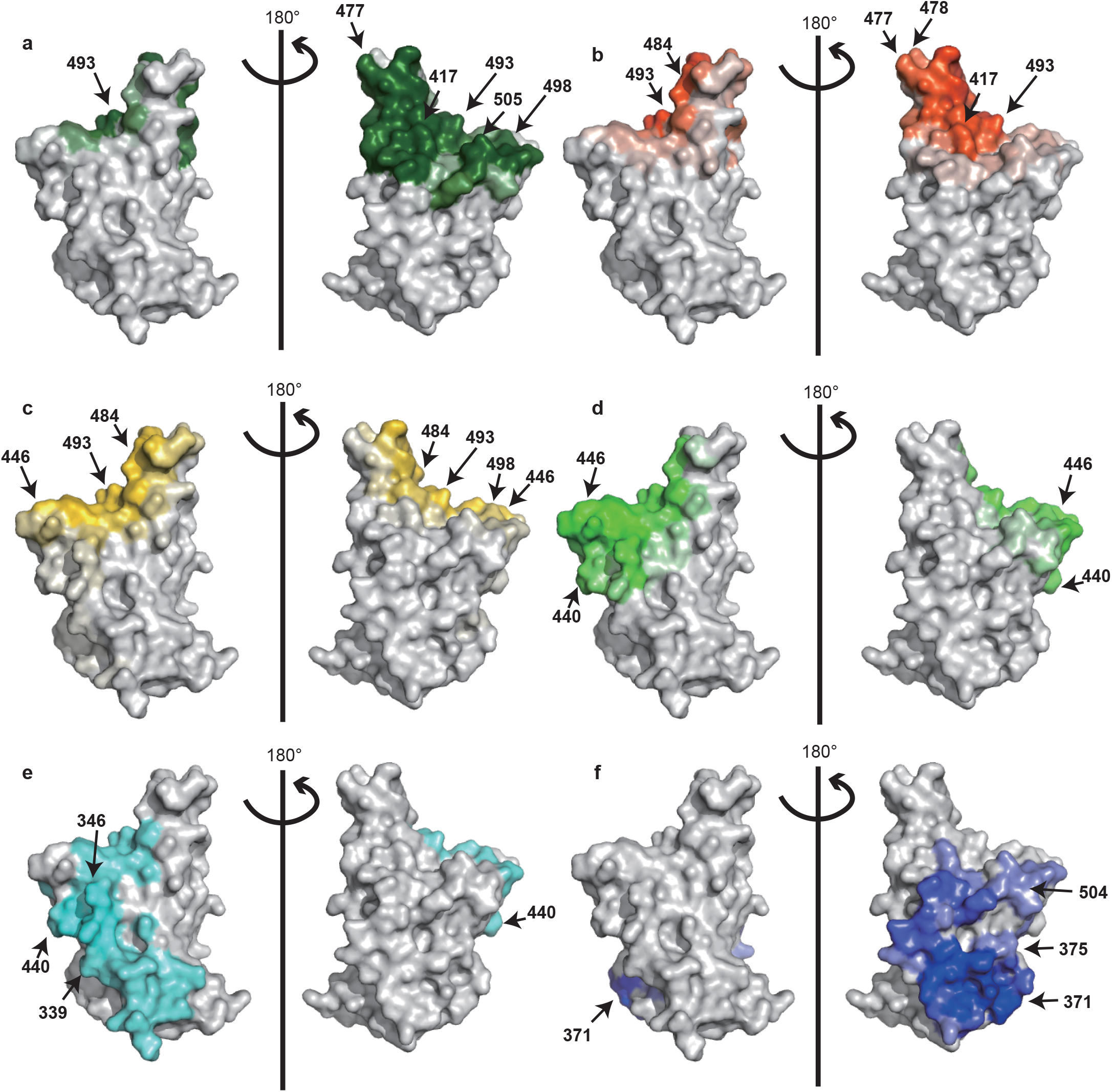
Antibody-RBD interface distribution for NAbs of each epitope group. **a-f**, Aggregated antibody-antigen interface of antibodies for epitope group A-F, respectively. Antibody-antigen interface was indicated from publicly available structures of neutralizing antibodies in complex with SARS-CoV-2 RBD. Different colors distinguish epitope groups, and the shade reflects group-specific site popularity to appear on the complex interface. Shared interface residues (Omicron) of each group are annotated.

**Extended Data Fig. 8:**
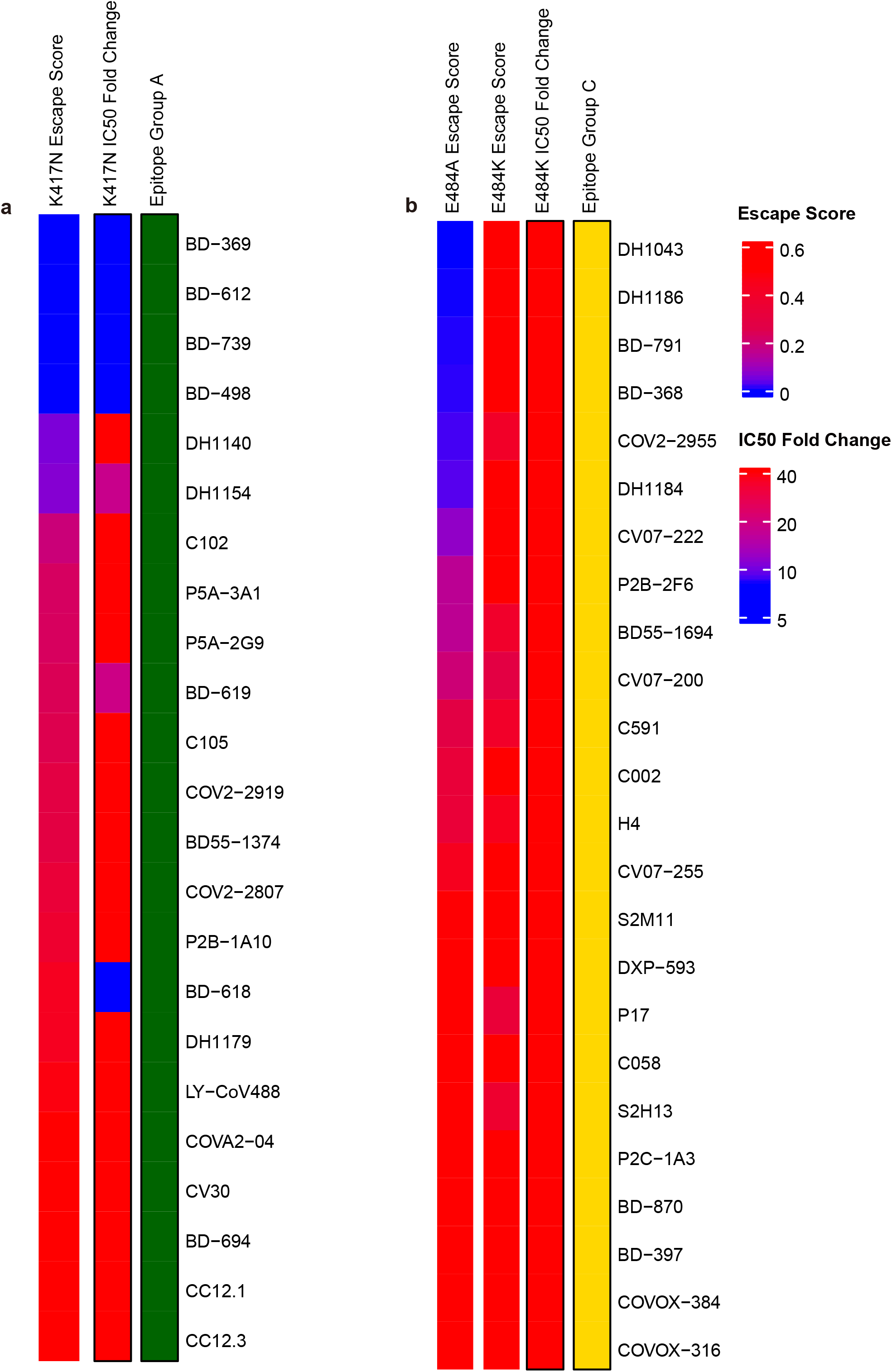
Comparison between mutation escape scores estimated from yeast display and neutralization of variants carrying corresponding mutations. **a**, K417N escape scores and corresponding K417N pseudovirus neutralizing IC50 fold change compared to D614G pseudovirus of antibodies within epitope group A. **b**, E484K/E484A escape scores and corresponding E484K pseudovirus neutralizing IC50 fold change compared to D614G pseudovirus of antibodies within epitope group C.

**Extended Data Fig. 9:**
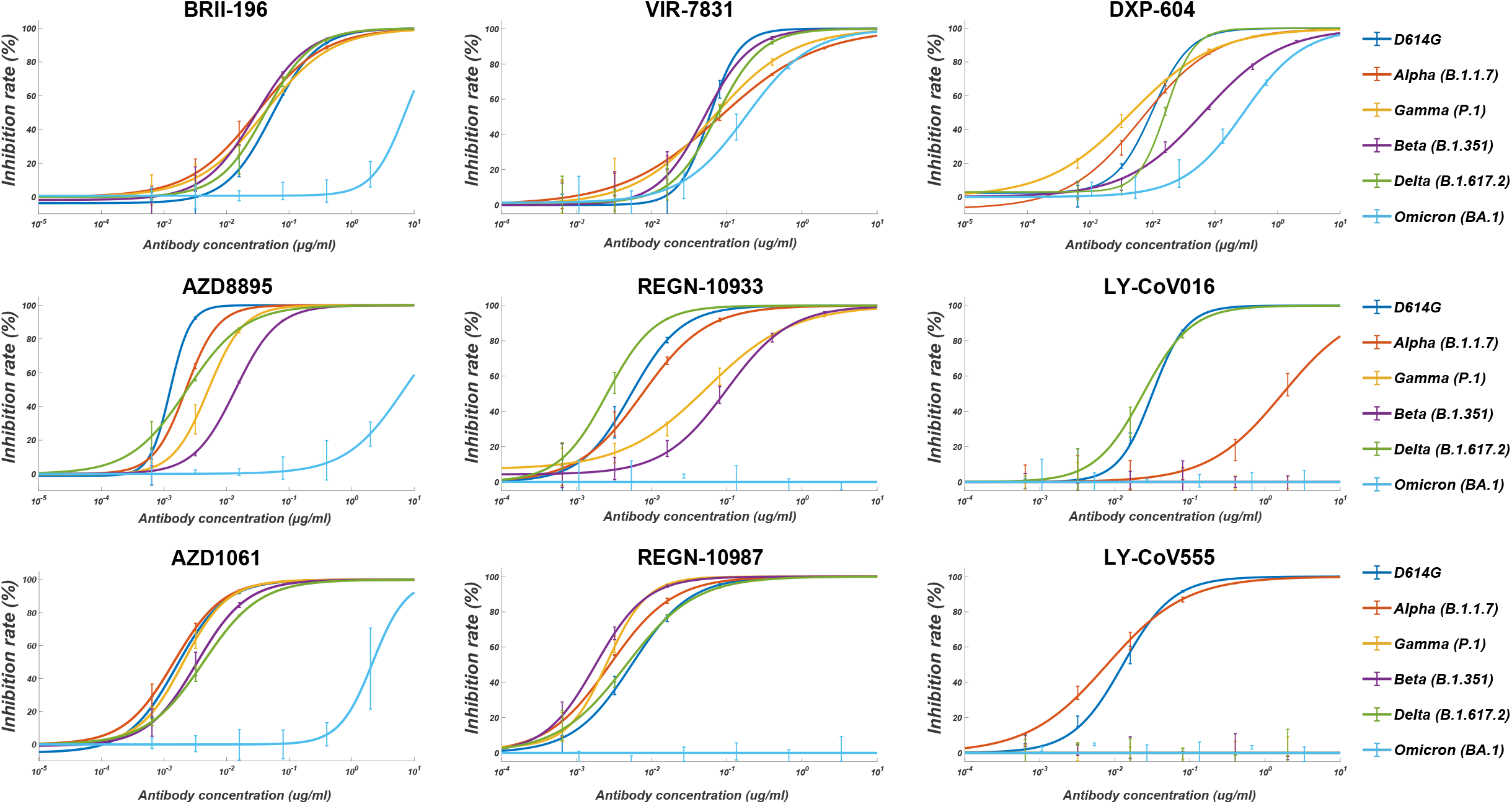
Pseudovirus neutralization of NAb drugs against SARS-CoV-2 variants of concern. Pseudovirus (VSV-based) assays were performed using Huh-7 cells. Data are collected from three biological replicates and represented as mean±s.d.

**Extended Data Fig. 10:**
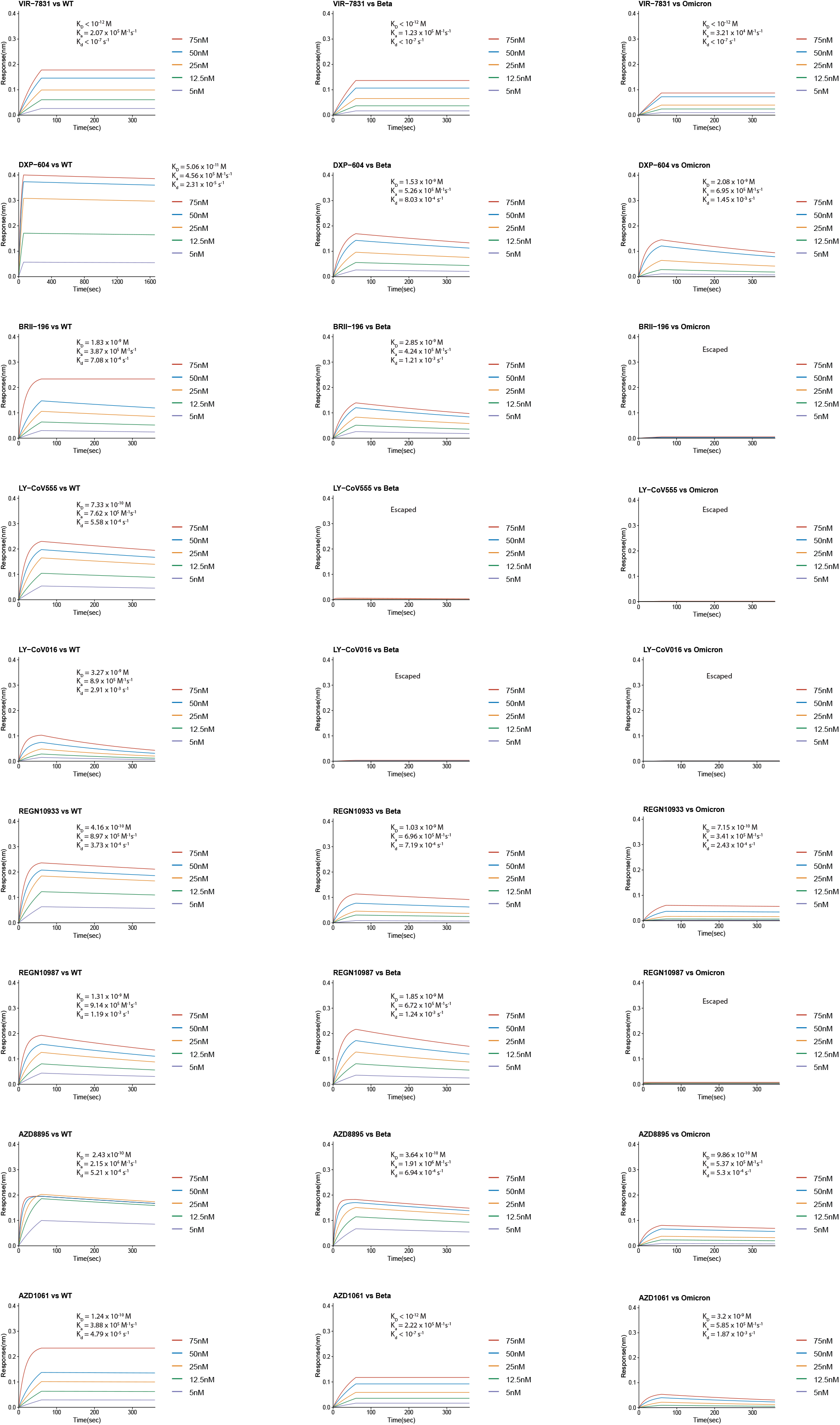
BLI response between NAb drugs and the RBD of SARS-CoV-2 wildtype, Beta, or Omicron strain. Antibodies were captured by Protein A sensor. The concentrations of RBD are shown in different colors. Dissociation constant (K_D_), association constant (ka), and dissociation rate constant (kd) are labeled. NAbs without binding are marked as “Escaped”.

