## Supplementary Data 1 for "Omicron escapes the majority of existing SARS-CoV-2 neutralizing antibodies"

### Escape maps of Epitope Group A antibodies

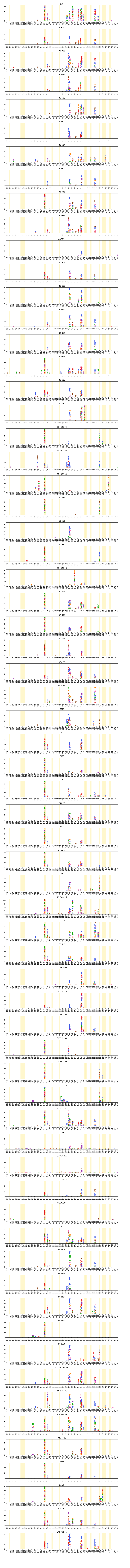

### Escape maps of Epitope Group B antibodies

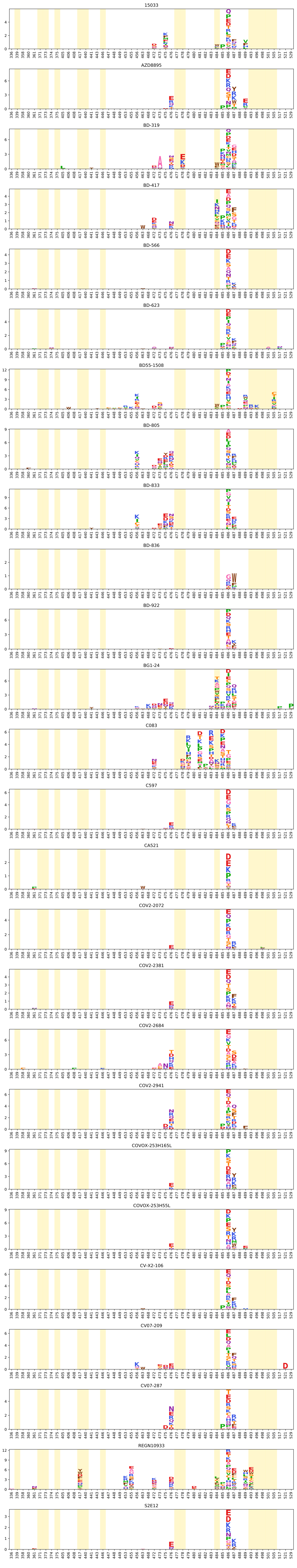

### Escape maps of Epitope Group C antibodies

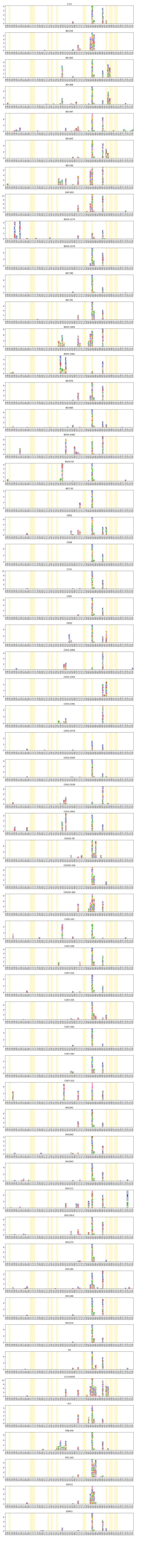

**Escape maps of Epitope Group D antibodies**

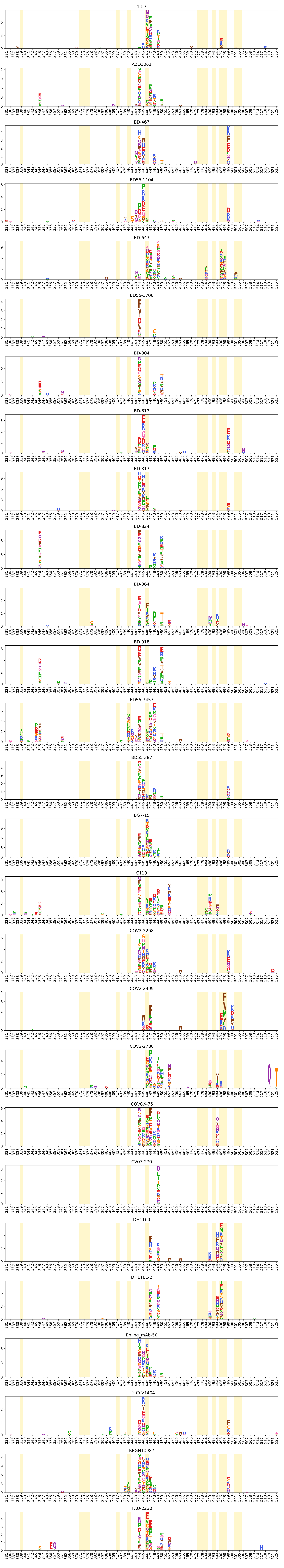

**Escape maps of Epitope Group E antibodies**

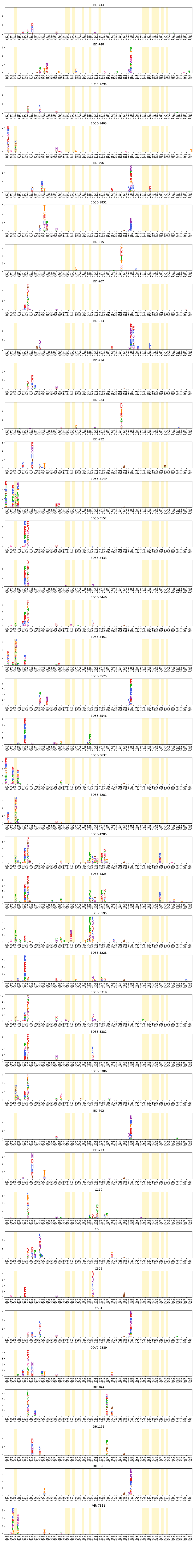

### Escape maps of Epitope Group F antibodies

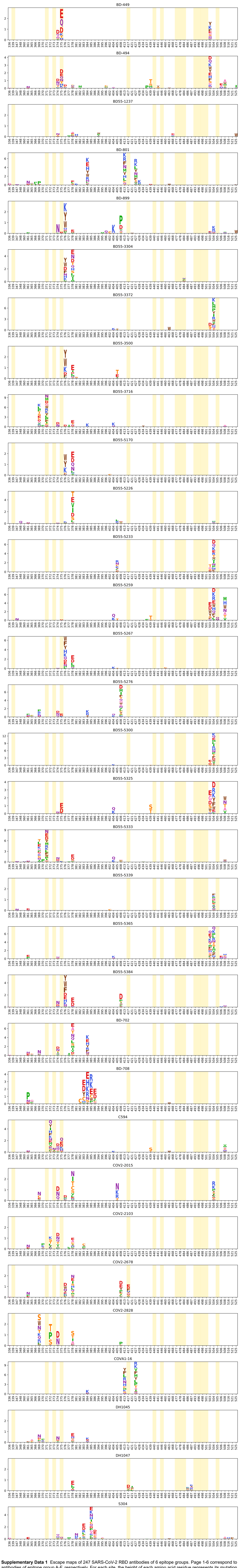


**Supplementary Data 1** Escape maps of 247 SARS-CoV-2 RBD antibodies of 6 epitope groups. Page 1-6 correspond to antibodies of epitope group A-F, respectively. For each site, the height of each amino acid residue represents its mutation escape score. Sites mutated frequently in Omicron variant are highlighted. Residues are colored according to their functional groups (*dmslogo_funcgroup* option of *logomaker* Python package).
