## Supplementary Data 2 for "Omicron escapes the majority of existing SARS-CoV-2 neutralizing antibodies"

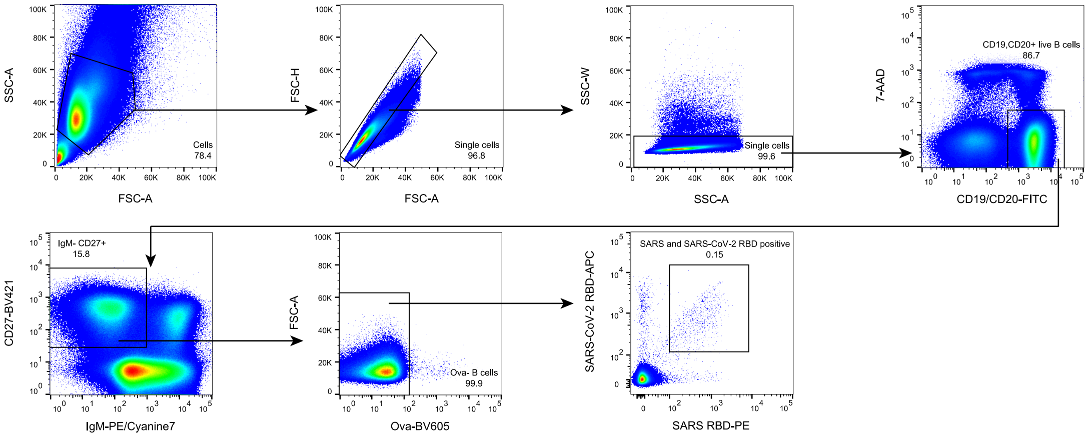


**Supplementary Data 2** Flow cytometry gating scheme for antigen-specific memory B cell sorting. Gating was on singlets that were 7-AAD$^-$,CD19 or CD20$^+$, IgM$^-$, CD27$^+$, and Ova$^-$. Sorted cells were SARS RBD$^+$ and SARS-CoV-2 RBD$^+$.
